# A partially shared joint clustering framework for detecting protein complexes from multiple state-specific signed interaction networks

**DOI:** 10.1101/2023.01.16.524205

**Authors:** Youlin Zhan, Jiahan Liu, Min Wu, Chris Soon Heng Tan, Xiaoli Li, Le Ou-Yang

## Abstract

Detecting protein complexes is critical for studying cellular organizations and functions. The accumulation of protein-protein interaction (PPI) data enables the identification of protein complexes computationally. Although various computational approaches have been proposed to detect protein complexes from PPI networks, most of them ignore the signs of PPIs that reflect the ways proteins interact (activation or inhibition). As not all PPIs imply cocomplex relationships, taking into account the signs of PPIs can benefit the detection of protein complexes. Moreover, PPI networks are not static, but vary with the change of cell states or environments. However, existing protein complex identification algorithms are primarily designed for single-network clustering, and rarely consider joint clustering of multiple PPI networks. In this study, we propose a novel partially shared signed network clustering model (PS-SNC) for detecting protein complexes from multiple state-specific signed PPI networks jointly. PS-SNC can not only consider the signs of PPIs, but also identify the common and unique protein complexes in different states. Experimental results on synthetic and real datasets show that PS-SNC outperforms other state-of-the-art protein complex detection methods. Extensive analysis on real datasets demonstrate the effectiveness of PS-SNC in revealing novel insights about the underlying patterns of different cell lines.

## 1. Introduction

Proteins are key executors of almost all cellular processes and often perform their specific biological functions through interactions with other proteins to form protein complexes. Disruptions or dysregulation of protein complexes often lead to cell dysfunction that may manifest as disease [1, 2]. Therefore, the identification and analysis of protein complexes is crucial not only for understanding the mechanisms behind the functional organization of cells but also the pathogenesis of diseases, which could provide insights for disease diagnosis and drug development [3, 4]. Various biological experimental techniques, such as tandem affinity purification-mass spectrometry (AP-MS) [5, 6] and yeast-two hybrid (Y2H) [7, 8], have been adapted for large-scale identification and study of protein complexes. However, Y2H does not detect protein complexes directly but identify interaction between two proteins while AP-MS often detect subsets of different protein complexes [9]. With the development of high-throughput protein interaction profiling technologies, a large amount of protein-protein interaction (PPI) data has been accumulated, enabling the identification of protein complexes from PPI networks using computational methods instead [10]. Recently, a great number of computational efficient methods have been proposed to identify protein complexes from PPI networks [11, 12, 13, 14, 15, 16, 17, 18, 19, 20, 21, 22]. Most of these methods are based on the assumption that interacting protein pairs tend to belong to same complexes and densely connected subnetworks in PPI networks are potential protein complexes.

Most existing protein complex detection methods do not consider the signs of PPIs (i.e., activation-inhibition relationships). Studies have shown that proteins belonging to same complexes are mainly connected by positive interactions, while negative interactions are more likely to occur between proteins belonging to different complexes [23]. Recently, Huttlin *et al*. [24] generated the most complete dataset of the human interactome to date, named BioPlex 3.0, which includes two PPI networks obtained in HEK293T cells and HCT116 cells that were derived from embryonic kidney tissue and colorectal carcinoma respectively. They calculated the correlation of interacting protein pairs in the BioPlex networks, resulting in sign information of protein interactions, and found that positively correlated proteins (i.e., positive PPIs) tend to belong to same complexes, while negatively correlated proteins (i.e., negative PPIs) were more likely to belong to different complexes. As positive and negative correlations imply different functional and structural relationships between proteins, considering the signs of PPIs can help to improve the accuracy of protein complex identification and deepen our understanding of the mechanism of cellular function. Ou-Yang *et al*. [25] proposed a signed network clustering model named SGNMF to identify protein complexes from a single signed PPI network, and confirmed that considering the signs of PPIs can indeed improve the accuracy of protein complex identification.

The above methods detect protein complexes by single-network clustering or multi-view network clustering [26, 27]. However, they seldom consider the changes of protein complexes in different states. In fact, the interactions between proteins are not static, but varies with the change in cell states or environments [24]. Protein complexes may also dynamically assemble or dissociate as needed [28]. On one hand, core protein complexes that serve as the backbone of cellular activities are often relatively stable but on another hand, some function-specific protein complexes may be formed only under specific conditions [24]. For example, the ideal targets for cancer therapy should be found in most cancer cells, instead of normal cells. Thus, complexes as ideal targets are those formed significantly different in different tumor types [29]. Therefore, instead of analyzing each PPI network separately, we need to analyze multiple PPI networks that under different cell states jointly to detect the common protein complexes that are shared across different states as well as identifying protein complexes that uniquely exist in certain states. Adopting this strategy could improve the accuracy of protein complex identification through joint analysis while at the same time helps to reveal various protein complexes underlying various cell state transitions and adaptations.

To address the above challenges, in this study, we develop a novel partially shared signed network clustering (PS-SNC) model to identify the common and unique protein complexes in two state-specific signed PPI networks simultaneously. The overall framework of our model is shown in Fig. 1. Firstly, we introduce a partially shared non-negative matrix factorization model to identify protein complexes in two state-specific signed PPI networks jointly, and divide the predicted complexes of each network into two parts, i.e., the common complexes that are shared across two networks and the unique complexes that are specific to each network. Secondly, to consider the sign information of PPIs when detecting protein complexes, we introduce a signed graph regularization term. Furthermore, we introduce Hilbert-Schmidt Independence Criterion (HSIC) as a diversity constraint to penalize the correlations between network-specific parts, and a low-rank constraint is employed to control the number of generated clusters and the overlap among clusters. Extensive experimental results on synthetic and real datasets show the superiority of our proposed method over five state-of-the-art protein complex detection methods.

**Fig. 1.**
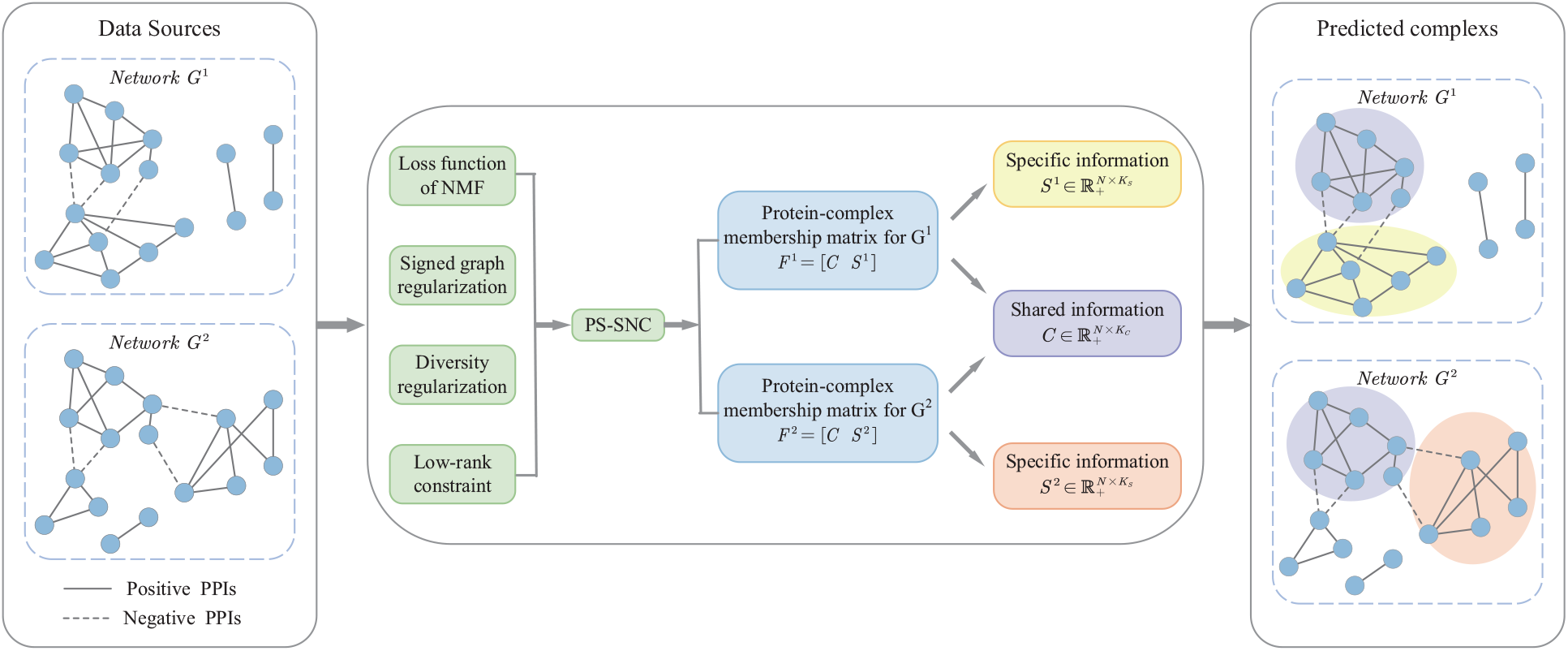
The overall framework of PS-SNC. Given two PPI networks with positive edges and negative edges, our PS-SNC can detect the common protein complexes that are shared across two networks (the complex with purple color) and the unique complexes that are specific to each network (a complex with yellow color for network *G*1 and a complex with pink color for network *G*^2^). In particular, PS-SNC is a NMF framework with 3 additional regularization terms.

## 2. Method

In this section, we describe the details of our proposed Partially Shared Signed Network Clustering (PS-SNC) model.

### 2.1. Model formulation

Given two signed PPI networks *G*^1^ and *G*^2^, let *V* ^*m*^ denotes the set of proteins in *G*^*m*^ for *m* = 1, 2, and *N* denotes the total number of proteins in both networks, i.e., *N* = | *V* ^1^ ∪ *V* ^2^ |. Each signed network is described by an adjacency matrix *A*^*m*^ ∈ ℝ^*N*×*N*^, where 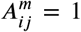 means there is a positive PPI between *i*-th and *j*-th proteins, 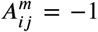 means there is a negative PPI between *i*-th and *j*-th proteins, and 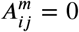 means there is no known interaction between *i*-th and *j*-th proteins. Moreover, we divide *A*^*m*^ into two parts, i.e., *A*^*m*^ = *A*^*m*+^ − *A*^*m*−^, where *A*^*m*+^ and *A*^*m*−^ denote the positive and negative parts of *A*^*m*^ respectively. Here, 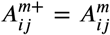 if 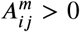, and 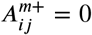 otherwise. 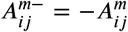 if 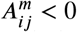, and 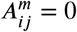 otherwise. Our goal is to identify protein complexes from these two signed PPI networks jointly, and identify the common complexes that are shared across two networks and the unique complexes that are specific to each network simultaneously.

As positive interactions usually take place between proteins belong to same complexes [23], the elements in the positive adjacency matrix *A*^*m*+^ describe the co-complex relationships between proteins. Thus, we first introduce a nonnegative indication matrix 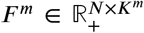 to mine the underlying co-complex relationships between proteins by approximating *A*^*m*+^ as follows:

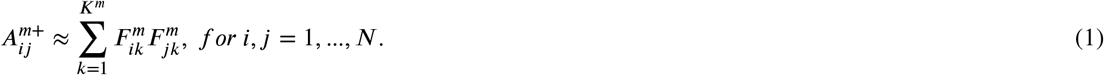

Here, *K*^*m*^ denotes the number of complexes in *m*-th network and each element 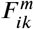 of *F* ^*m*^ describes the propensity of *i*-th protein to belong to *k*-th complex. A higher value of 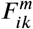 indicates a higher propensity of *i*-th protein to belong to *k*-th complex. As we allow a protein to have high propensities on more than one complex, our model supports the identification of overlapping protein complexes.

To measure the distance between *F* ^*m*^(*F* ^*m*^)^*T*^ and *A*^*m*+^, following previous studies [30], we adopt Kullback-Leibler (KL) divergence. Since different networks may cover different numbers of proteins, for each network, we introduce a vector *θ*^*m*^ ∈ {0, 1}^*N*×1^ to indicate the proteins belong to each network, where 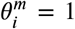 means that *i*-th protein is included in network *A*^*m*+^, and 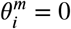 otherwise. The loss function is defined as follows:

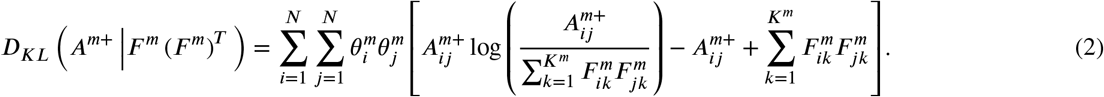

Instead of exploring the consistent patterns of different networks and forcing *F* ^1^ = *F* ^2^, we try to identify the common and unique complexes in different networks jointly. Thus, in this study, *F* ^*m*^ is divided into two parts, i.e., *F* ^*m*^ = [*C, S*^*m*^], 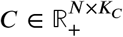 where is the common part that reflects the consistent information shared across the two networks and 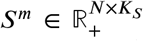 is the unique part that reflects the specific information of each network, and *K*_*C*_ and *K*_*S*_ are the dimensions of the common and unique latent factors respectively. Similar to the choice of [31], we set *K*_*C*_ : *K*_*S*_ = 2 : 1 in our experiments. Accordingly, the above loss function can be modified as follows:

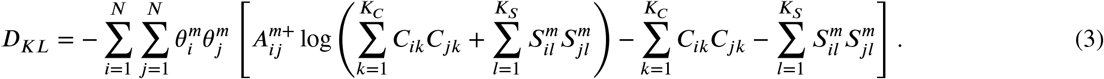

### 2.2. Signed graph regularization

As positive PPIs usually take place between proteins belonging to same complexes and negative PPIs are more likely to be inter-module interactions [32], we need to take into account the signs of PPIs when detecting protein complexes. Graph Laplacian regularizer is widely used to measure the smoothness of latent representations of nodes based on their similarities [33]. In this study, considering the signs of PPIs, we introduce a signed graph Laplacian term to regularize the propensities of interacting proteins on the same complexes [25]. As positive PPIs indicate co-complex relationships and negative PPIs indicate inter-module relationships, the signed Laplacian regularization for *F* ^*m*^ is defined as follows:

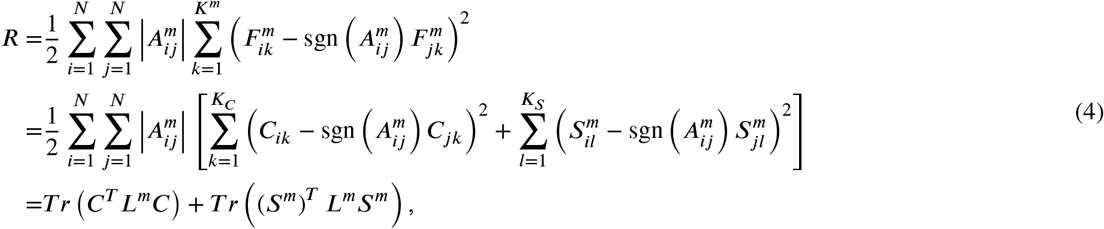

where *T r*(·) represents the trace of the matrix and 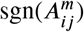 denotes the sign of 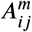.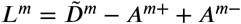 is the Laplace matrix, where 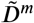 is the diagonal matrix defined by 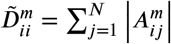.

### 2.3. Diversity regularization

In order to explore the unique complexes specific to certain networks, we introduce a diversity constraint to measure the difference between unique parts *S*^1^ and *S*^2^. In particular, we employ the Hilbert-Schmidt Independence Criterion (HSIC) to construct the diversity constraint. HSIC can measure the dependence of variables by mapping variables to a Reproducing Kernel Hilbert Space (RKHS), which can measure more complicated correlations. Moreover, HSIC is computationally efficient as it does not need to estimate the joint distribution of random variables explicitly [34]. Therefore, we adopt HSIC to penalize the correlation between *S*^1^ and *S*^2^, encouraging the identification of unique complexes. Here, we use an inner product kernel for HSIC, and the estimator of *HSIC*(*S*^1^, *S*^2^) is given as follows [35]:

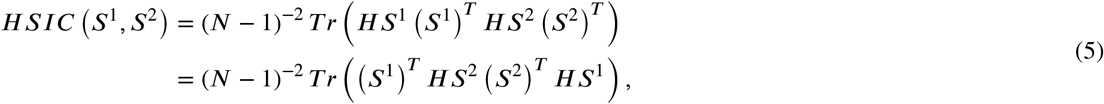

where *H* = *I* − 1/*N*, and *I* is the N-order identity matrix.

### 2.4. Low-rank constraint

Note that *F* ^*m*^(*F* ^*m*^)^*T*^ represents the potential co-complex propensities between proteins, its rank should, ideally, be equal to the number of complexes. As we have no prior knowledge about the number of complexes in each network, we introduce a low-rank constraint on *F* ^*m*^(*F* ^*m*^)^*T*^ such that our model could determine the number of clusters adaptively. We adopt the trace norm constraint ‖ *F* ^*m*^ (*F* ^*m*^)^*T*^‖_*_, which is a relaxation of the low-rank constraint [26], to achieve this goal. According to the above definition, we have 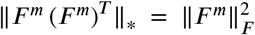, where ‖·‖_*F*_ denotes the Frobenius norm.

### 2.5. Objective function

Considering the above factors, the overall objective function of our PS-SNC model can be expressed as:

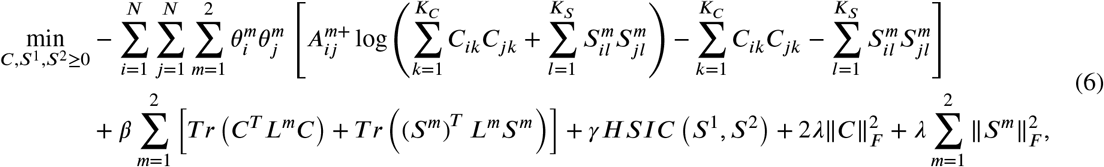

where *β, γ* and *λ* control the trade-off between smoothness, diversity and low-rank constraints, respectively.

#### Algorithm 1

**Algorithm for PS-SNC**

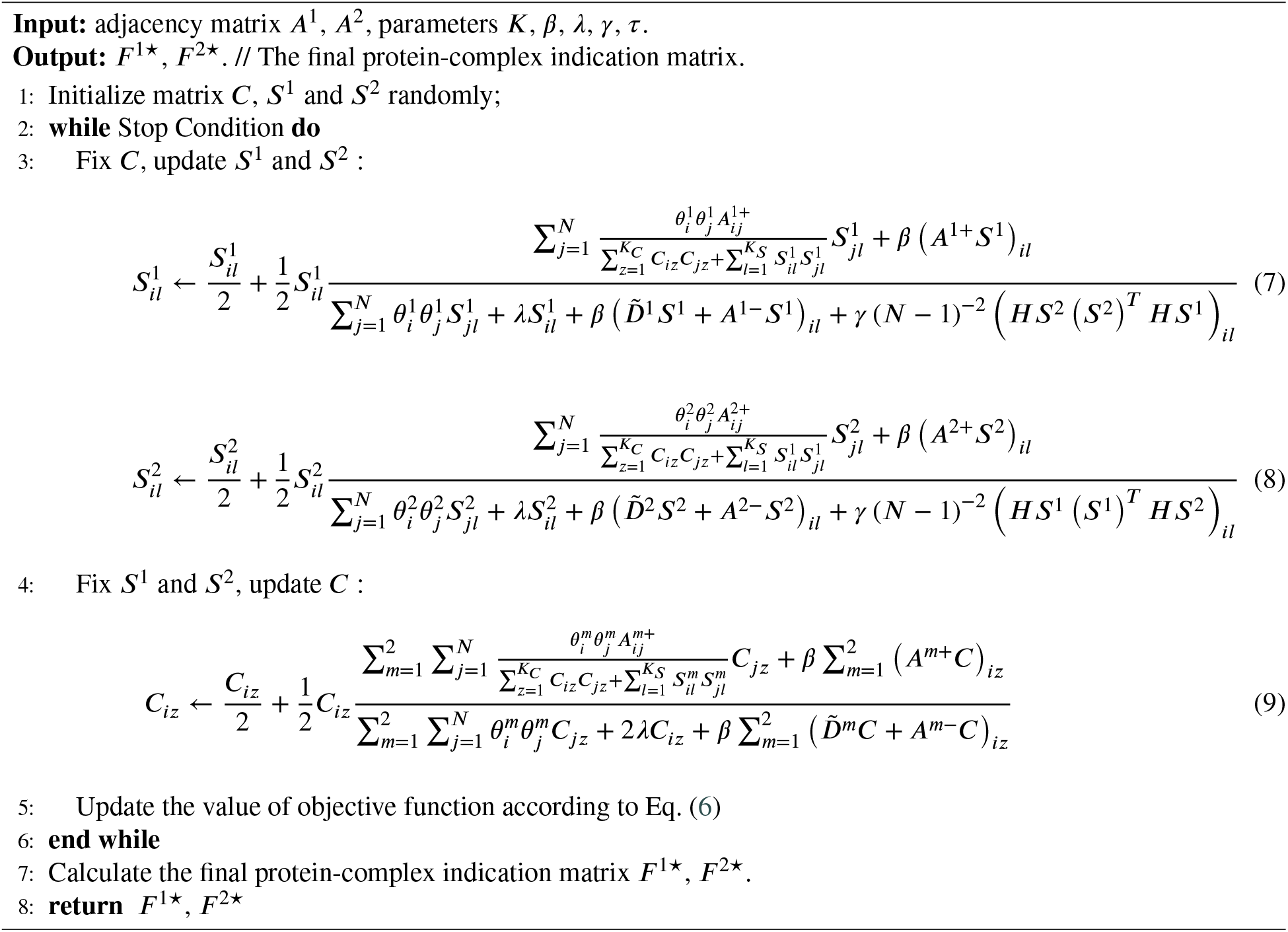

### 2.6. Parameter estimation

As the objective function is not a joint convex function over all variables *C, S*^1^, and *S*^2^, we utilize an alternating optimization strategy to solve the optimization problem in Eq. (6). Specifically, we optimize one variable of the objective function each time while fixing other variables. According to the multiplicative update rule [30, 36], we can get the update rules for *C, S*^1^, and *S*^2^ as shown in Algorithm 1.

Given the initial values of *C, S*^1^, and *S*^2^, we update *C, S*^1^, and *S*^2^ iteratively, until the stopping criterion is satisfied. In this study, we stop the iterations until the relative change of the objective function is less than 1e-6 or the number of iterations reaches a predefined maximum value, which is set to 200. As the objective function in lem in Eq. (6) is non-convex, updating *C, S*^1^, and *S*^2^ according to the update rule may converge to a local optimum, and the estimators of *C, S*^1^, and *S*^2^ rely on their initial values. In order to reduce the risk of local minimum, the entire updating process is repeated 10 times with random restarts and the minimizer of the objective function are treated as the final estimators of *C, S*^1^, and *S*^2^, denoted as *Ĉ, Ŝ*^*1*^, *Ŝ*^2^, respectively.

Since the elements in 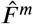 are all continuous values, describing the propensity of the *i*-th protein to belong to the *k*-th predicted complex. Following previous studies [25], we use the following rules to discretize 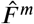 and obtain the final protein-complex indication matrix *F* ^*m*⋆^.

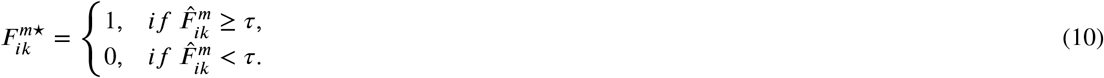

Here, 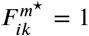 means the *i*-th protein is belong to the *k*-th predicted complex. In the experiments, we found that *τ* = 0.1 can always get reasonable results, so we fix *τ* = 0.1 in this study. Algorithm 1 summarizes the implementation details of our PS-SNC model. The computational complexity of updating *C* and *S*^*m*^ once is *O*(*N*^2^*K*_*C*_) and *O*(*N*^2^*K*_*S*_), respectively. If the number of iterations is the predefined maximum value *Iter*, the total time cost of PS-SNC is *O*(*Iter*(*N*^2^*K*_*C*_ + 2*N*^2^*K*_*S*_)). Considering that the true PPI networks are usually very sparse, the overall computational cost is *O*(*Iter*(| *E*^+^ | + | *E*^−^|)(*K*_*C*_ + 2*K*_*S*_)), where *E*|^+^ | and | *E*^−^| denote the number of positive and negative PPIs, respectively.

## 3. Experiments

In this section, we demonstrate the advantages of PS-SNC through experiments on both synthetic and real datasets.

### 3.1. Experimental settings

#### 3.1.1. Evaluation metrics

We adopt three evaluation metrics to comprehensively evaluate the performance of various methods, i.e., accuracy (ACC) [37], F-measure, and fraction of matching complexes (FRAC) [16].

ACC is defined as the geometric mean of sensitivity (Sn) and positive predictive value (PPV). Let *B*_*ij*_ denote the number of proteins shared between true complex *t*_*i*_ and predicted complex *p*_*j*_. Sn, PPV and ACC are defined as follows:

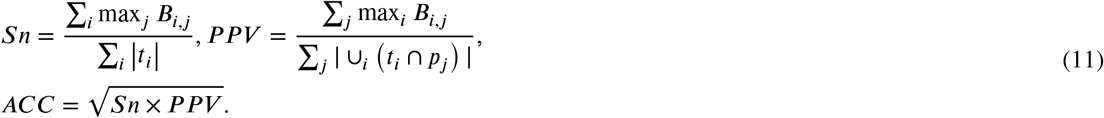

Given a true complex *t*_*i*_ and a predicted complex *p*_*j*_, the overlap fraction between them is defined as:

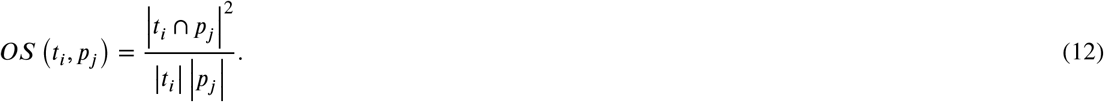

According to previous studies [12, 38], we consider two complexes to be matched if the overlap score between them is greater than or equal to 0.2. Let TP (true positive) be the number of predicted complexes that are matched by the true complexes, and FN (false negative) be the number of true complexes that are not matched by the predicted complexes, and FP (false positive) be the number of predicted complexes minus TP. Precision, Recall and F-measure are defined as follows:

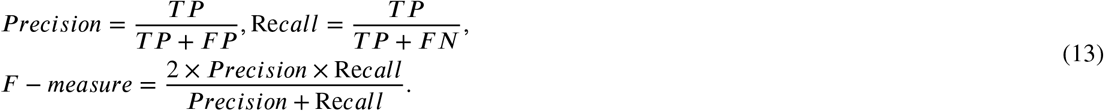

FRAC measures what fraction of complexes in the benchmark are matched by at least one predicted complex. FRAC is defined as follows:

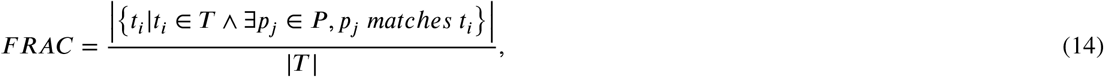

where *T* is the set of true complexes and *P* is the set of predicted complexes.

#### 3.1.2. Baselines

To evaluate the performance of PS-SNC, we compare PS-SNC with 5 state-of-the-art protein complex identification methods, including ClusterONE [16], IPCA [13], NCMine [21], PSMVC [26] and SGNMF [25]. Among these baseline methods, only SGNMF could take into account the signs of PPIs. Hence, we apply SGNMF and our PS-SNC on signed PPI networks. Meanwhile, we apply the remaining four methods, i.e., ClusterONE, IPCA, NCMine and PSMVC, on unsigned PPI networks, where the signs of interactions are ignored. Furthermore, since the four methods ClusterONE, IPCA, NCMine and SGNMF are designed to identify complexes from a single PPI network, we apply them on each PPI network separately, and evaluate the performance of these algorithms on the two networks respectively. For all algorithms, we discard their predicted complexes with less than three proteins (i.e., interactions only). Optimal parameters are set for IPCA, NCMine, PSMVC, SGNMF to obtain their best performance, while ClusterONE used the default parameters set by the authors.

### 3.2. Simulation studies

### 3.2.1. Synthetic datasets

We evaluate the performance of various algorithms on synthetic datasets to validate the benefits of joint clustering of multiple networks and considering the signs of PPIs. We generate two synthetic signed networks as follows. First, 3000 nodes are generated and then divided into 100 clusters with equal size, i.e, each cluster has 30 nodes. Among them, 50 clusters are shared to both networks. For another 20 clusters, each cluster is split into two sub-clusters with 10 overlapping nodes, which are considered as two partially shared clusters from two networks respectively. As such, each network has 20 partially shared clusters. For the remaining 30 clusters, we randomly select half of them for each network as network-specific clusters, i.e., each network has 15 network-specific clusters. Finally, each network has 85 clusters in total.

Then we generate edges in each of the two networks. Nodes within the same partially shared cluster or network-specific cluster have a probability *p*_*pos*_ = 0.15 to form a positive edge. Meanwhile, to further take noises into consideration, the signs of edges have a probability *p*_*noise*_ = 0.02 to be flipped. To demonstrate the benefit of joint clustering of multiple networks, we enforce the information contained in the shared clusters of the two networks more complementary. In network 1, nodes within the first 25 shared clusters have a probability *p*_*c*1_ to form a positive edge and are flipped with a probability *p*_*noise*1_ = 0.02. While nodes within the last 25 shared clusters have a probability *p*_*c*2_ = 0.05 to form a positive edge and are flipped with a probability *p*_*noise*2_ = 0.05. And it is the opposite in network 2. In addition, nodes within different clusters will have a probability *p*_*neg*_ to form negative edges and be flipped with probability *p*_*noise*_ = 0.02. And the value of *p*_*neg*_ varies from {0.0005, 0.005}, generating two types of networks: balanced networks and unbalanced networks. The value of *p*_*c*1_ varies from {0.1, 0.2, 0.3} to adjust the density of edges within the shared clusters of the two networks. Finally, a total of 6 datasets are generated. All the 6 datasets are available via https://github.com/Zyl-SZU/PS-SNC.

We evaluate the performance of various algorithms on each network separately, using the ground truth clusters of each network as the gold standard.

#### 3.2.2. Results on synthetic datasets

There are four parameters in our model: *K, β, γ*, and *λ. K* is the number of possible complexes, where *K* = *K*_*C*_ + 2*K*_*S*_. *β, γ*, and *λ* control the effects of the signed graph regularization term, diversity regularization term and low-rank constraint, respectively.

We find from the experiment results that *β* = 4 always achieves competitive results, so we fixed *β* = 4 in the following experiments. We perform grid search for *K* from {100, 200, 300}, for *λ* from {2^0^, 2^1^, …, 2^6^} and for *γ* from {0, *N*^2^ × 2^−1^, *N*^2^ × 2^0^, …, *N*^2^ × 2^4^} to obtain the best performance.

As show in Table 1, Datasets # 1, 2, and 3 are balanced networks with different density *p*_*c*1_, while Datasets # 4, 5, and 6 are unbalanced networks with different density *p*_*c*1_. We have the following observations. On all the synthetic networks, PS-SNC and SGNMF significantly outperform PSMVC, NCMine, IPCA and ClusterONE. Especially in unbalanced networks with a large number of negative edges, the performance of PSMVC, NCMine, IPCA and ClusterONE is greatly degraded due to the interference of negative edges. As both PS-SNC and SGNMF take into account the signs of interactions, the above results show that distinguishing between negative and positive interactions facilitates more accurate identification of protein complexes.

**Table 1.** The results of different methods on synthetic datasets

| Datasets | Methods | network # 1 |  |  |  |  | network # 2 |  |  |  |  |
| --- | --- | --- | --- | --- | --- | --- | --- | --- | --- | --- | --- |
|  |  | # complexes | # proteins | ACC | F-measure | FRAC | # complexes | # proteins | ACC | F-measure | FRAC |
| Dataset # 1 | PS-SNC | 102 | 1517 | **0.711** | **0.849** | **0.765** | 102 | 1515 | **0.711** | **0.833** | **0.765** |
|  | SGNMF | 292 | 1763 | 0.574 | 0.817 | 0.753 | 294 | 1771 | 0.570 | 0.804 | **0.765** |
|  | PSMVC | 140 | 2966 | 0.649 | 0.710 | **0.765** | 143 | 2966 | 0.649 | 0.701 | **0.765** |
|  | NCMine | 257 | 586 | 0.327 | 0.332 | 0.306 | 198 | 489 | 0.315 | 0.290 | 0.282 |
|  | IPCA | 1017 | 2671 | 0.390 | 0.268 | 0.565 | 1002 | 2693 | 0.371 | 0.219 | 0.518 |
|  | ClusterOne | 161 | 600 | 0.309 | 0.204 | 0.200 | 145 | 533 | 0.290 | 0.196 | 0.212 |
| Dataset # 2 | PS-SNC | 96 | 1621 | **0.780** | **0.874** | **0.765** | 96 | 1611 | **0.802** | **0.890** | **0.765** |
|  | SGNMF | 296 | 1772 | 0.562 | 0.849 | **0.765** | 295 | 1778 | 0.557 | 0.850 | **0.765** |
|  | PSMVC | 107 | 2907 | 0.641 | 0.751 | 0.753 | 101 | 2936 | 0.620 | 0.708 | 0.729 |
|  | NCMine | 414 | 801 | 0.354 | 0.513 | 0.482 | 390 | 739 | 0.343 | 0.545 | 0.435 |
|  | IPCA | 1069 | 2663 | 0.464 | 0.469 | 0.647 | 1019 | 2679 | 0.445 | 0.343 | 0.635 |
|  | ClusterOne | 193 | 735 | 0.338 | 0.321 | 0.376 | 215 | 812 | 0.338 | 0.310 | 0.388 |
| Dataset # 3 | PS-SNC | 106 | 1654 | **0.807** | **0.890** | **0.765** | 109 | 1603 | **0.804** | **0.888** | **0.765** |
|  | SGNMF | 284 | 1751 | 0.627 | 0.861 | **0.765** | 286 | 1758 | 0.637 | 0.850 | **0.765** |
|  | PSMVC | 129 | 2956 | 0.670 | 0.763 | **0.765** | 120 | 2939 | 0.656 | 0.741 | 0.753 |
|  | NCMine | 609 | 842 | 0.372 | 0.771 | 0.424 | 637 | 882 | 0.381 | 0.782 | 0.412 |
|  | IPCA | 304 | 767 | 0.337 | 0.533 | 0.329 | 301 | 800 | 0.357 | 0.533 | 0.365 |
|  | ClusterOne | 201 | 842 | 0.366 | 0.416 | 0.412 | 228 | 933 | 0.374 | 0.337 | 0.376 |
| Dataset # 4 | PS-SNC | 63 | 1395 | **0.694** | **0.828** | **0.741** | 63 | 1417 | **0.704** | **0.814** | **0.729** |
|  | SGNMF | 200 | 2072 | 0.549 | 0.713 | **0.741** | 200 | 2091 | 0.551 | 0.686 | 0.718 |
|  | PSMVC | 225 | 2477 | 0.219 | 0.077 | 0.141 | 225 | 2489 | 0.169 | 0.039 | 0.071 |
|  | NCMine | 335 | 812 | 0.259 | 0.107 | 0.165 | 323 | 784 | 0.266 | 0.125 | 0.200 |
|  | IPCA | 210 | 591 | 0.230 | 0.054 | 0.082 | 213 | 597 | 0.217 | 0.040 | 0.071 |
|  | ClusterOne | 191 | 688 | 0.183 | 0.029 | 0.047 | 176 | 648 | 0.184 | 0.031 | 0.047 |
| Dataset # 5 | PS-SNC | 89 | 1529 | **0.761** | **0.876** | **0.765** | 90 | 1535 | **0.755** | **0.878** | **0.765** |
|  | SGNMF | 290 | 2023 | 0.596 | 0.728 | 0.729 | 290 | 2042 | 0.592 | 0.729 | 0.741 |
|  | PSMVC | 116 | 2970 | 0.309 | 0.289 | 0.341 | 118 | 2966 | 0.294 | 0.227 | 0.271 |
|  | NCMine | 424 | 931 | 0.298 | 0.282 | 0.247 | 476 | 977 | 0.313 | 0.336 | 0.318 |
|  | IPCA | 236 | 671 | 0.272 | 0.140 | 0.188 | 259 | 711 | 0.267 | 0.081 | 0.141 |
|  | ClusterOne | 191 | 743 | 0.204 | 0.065 | 0.094 | 204 | 754 | 0.208 | 0.055 | 0.094 |
| Dataset # 6 | PS-SNC | 94 | 1572 | **0.772** | **0.873** | **0.753** | 96 | 1544 | **0.769** | **0.851** | **0.765** |
|  | SGNMF | 69 | 969 | 0.636 | 0.688 | 0.576 | 84 | 989 | 0.638 | 0.736 | 0.635 |
|  | PSMVC | 75 | 2918 | 0.362 | 0.360 | 0.329 | 75 | 2920 | 0.342 | 0.313 | 0.294 |
|  | NCMine | 603 | 1046 | 0.340 | 0.568 | 0.329 | 640 | 1078 | 0.340 | 0.552 | 0.329 |
|  | IPCA | 329 | 866 | 0.308 | 0.338 | 0.306 | 292 | 808 | 0.307 | 0.314 | 0.259 |
|  | ClusterOne | 193 | 781 | 0.226 | 0.086 | 0.118 | 188 | 750 | 0.227 | 0.102 | 0.153 |

Datasets # 1, 2, and 3 are balanced networks with different density $p_{c1}$ . Datasets # 4, 5, and 6 are unbalanced networks with different density $p_{c1}$ . The best results for each network using each metric are in bold.

Among methods that do not consider the signs of interactions, PSMVC achieves the best performance in balanced networks, and has the same FRAC as PS-SNC and surpasses SGNMF in terms of ACC. Moreover, we can observe from Table 1 that the performance of PS-SNC is much more stable than SGNMF in terms of all evaluation metrics, demonstrating the benefits of joint clustering of multiple networks.

### 3.3. Real data analysis

#### 3.3.1. Real datasets

By considering the correlations of interacting protein pairs in the BioPlex networks as the signs of PPIs [24], we collected two signed networks, named Sign-293T and Sign-HCT116, to evaluate the performance of various protein complex identification methods. We summarize the statistics of the dataset in Table 2.

**Table 2.**
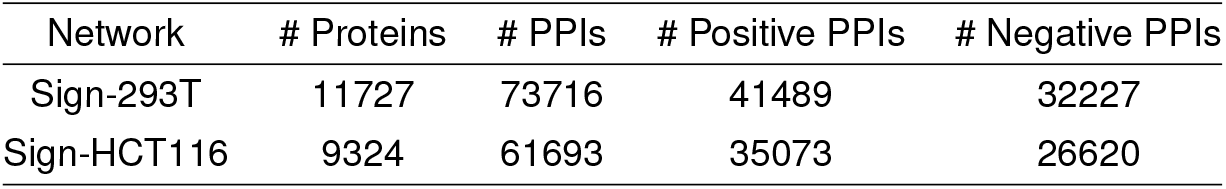
The statistics of real datasets

To measure whether the predicted complexes match known complexes, we employ the CORUM database [39] as the gold standards. To avoid selection bias, we filtered out proteins that are not involved in both PPI networks. Furthermore, we only considered complexes containing at least 3 or more proteins. Finally, CORUM contains 1774 complexes covering 2807 proteins. Since CORUM does not provide the state and cell line information of protein complexes, for single-network clustering methods, we merge their predicted complexes from the two networks. In particular, for predicted complexes *a* and *b*, if *OS*(*a, b*) ≥ 0.8, we consider them to be highly overlapping, and they are merged into one complex [38, 40].

#### 3.3.2. Results on real datasets

In this section, we present the experiment results of different methods with respect to CORUM on two real signed PPI networks: Sign-293T and Sign-HCT116. Since the reference complexes in CORUM are far from complete, the predicted complexes that do not match with any reference complexes are not necessarily undesired results. Instead, they may be potential protein complexes not covered by the reference set [16, 37]. Thus, following previous studies [26, 41, 27], in real data analysis, we do not use F-measure to evaluate the performance of various methods. The results are shown in Table 3. We can find from this table that PS-SNC outperforms other 5 methods in terms of ACC and FRAC. For instance, PS-SNC achieves ACC 0.656 and FRAC 0.378, which is 4.8% and 5% higher than the second best ACC and FRAC. FRAC can clearly indicate the effectiveness of algorithms in identifying reference complexes. The size and quality of predicted complexes is another important factor measured by ACC. Overall, the performance of PS-SNC on the real dataset is generally better than all the compared algorithms.

**Table 3.** The results of various methods on real datasets

| Methods | # complexes | # proteins | ACC | FRAC |
| --- | --- | --- | --- | --- |
| PS-SNC | 2351 | 8363 | **0.656** | **0.378** |
| SGNMF | 2842 | 9201 | 0.626 | 0.350 |
| PSMVC | 4128 | 10158 | 0.606 | 0.359 |
| NCMine | 2869 | 4363 | 0.559 | 0.346 |
| IPCA | 7241 | 9908 | 0.626 | 0.360 |
| ClusterOne | 760 | 3448 | 0.566 | 0.236 |

The best results for each network using each metric are in bold.

Furthermore, we calculate the proportion of positive PPIs within predicted complexes and reference complexes, and show the results of various methods in Fig. 2. As shown in this figure, the proportion of positive PPIs in most complexes predicted by PS-SNC and SGNMF are close to 1. Compared with the other four algorithms that do not consider the signs of PPIs, the predicted complex sets of PS-SNC and SGNMF have the same median and more similar data distributions as the reference complex set, indicating that considering the signs of PPIs can improve the quality of the predicted complexes by using the sign information to guide the clustering, while reducing the interference of negative PPIs.

**Fig. 2.**
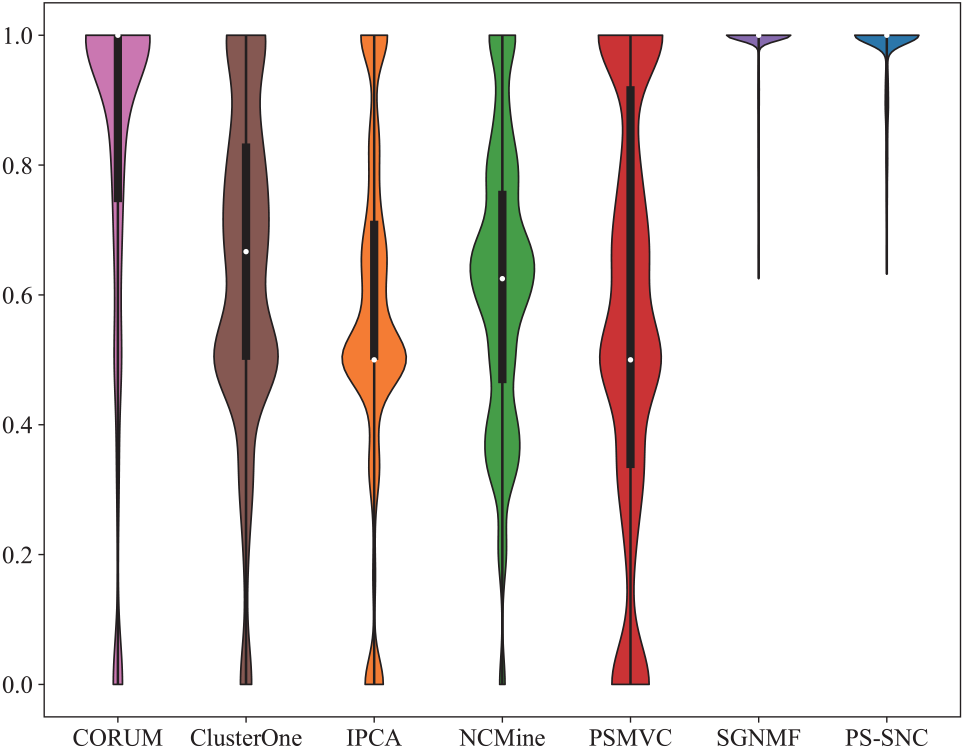
The proportion of positive PPIs per complex. The box in each violin plot shows the IQR and the median are highlighted by white dots.

#### 3.3.3. Parameter sensitivities

In this section, we investigate the parameter sensitivities of PS-SNC. We set the ranges of *λ* and *γ* according to the performance of PS-SNC on synthetic datasets, and the range of *K* according the scales of PPI networks. In particular, we first keep *K* = 12000, and run PS-SNC with different combination values of *λ*(*λ* ∈ {2^2^, 2^3^, …, 2^7^}) and *γ*(*γ* ∈ {0, *N*^2^ × 1, *N*^2^ × 2, …, *N*^2^ × 5}), and assess how well the predicted complexes match with CORUM reference set. Then we fix the values of *λ* and *γ* which result in the best performance, and study the effect of *K* on the performance of PS-SNC by setting *K* = 2000, 4000, …, 14000.

As shown in Fig. 3, for a fixed value of *γ*, as the value of *λ* increases, the ACC increases initially and decreases after reaching the maximum. Similarly, for a fixed value of *λ*, ACC first increases and then decreases as the value of *γ* increases. Thus, both *λ* and *γ* contribute to improve the performance of PS-SNC. Meanwhile, when *λ* < 2^7^, we observe that FRAC is not sensitive to the settings of *λ* and *γ*. On the other hand, we can find from Fig. 4 that with the increase of *K*, the FRAC increases initially and decreases after reaching the maximum, and ACC tend to be stable after increasing. Overall, PS-SNC achieves competitive performance when *K* = 12000, *λ* = 2^6^ and *γ* = *N*^2^ × 4 on the real dataset.

**Fig. 3.**
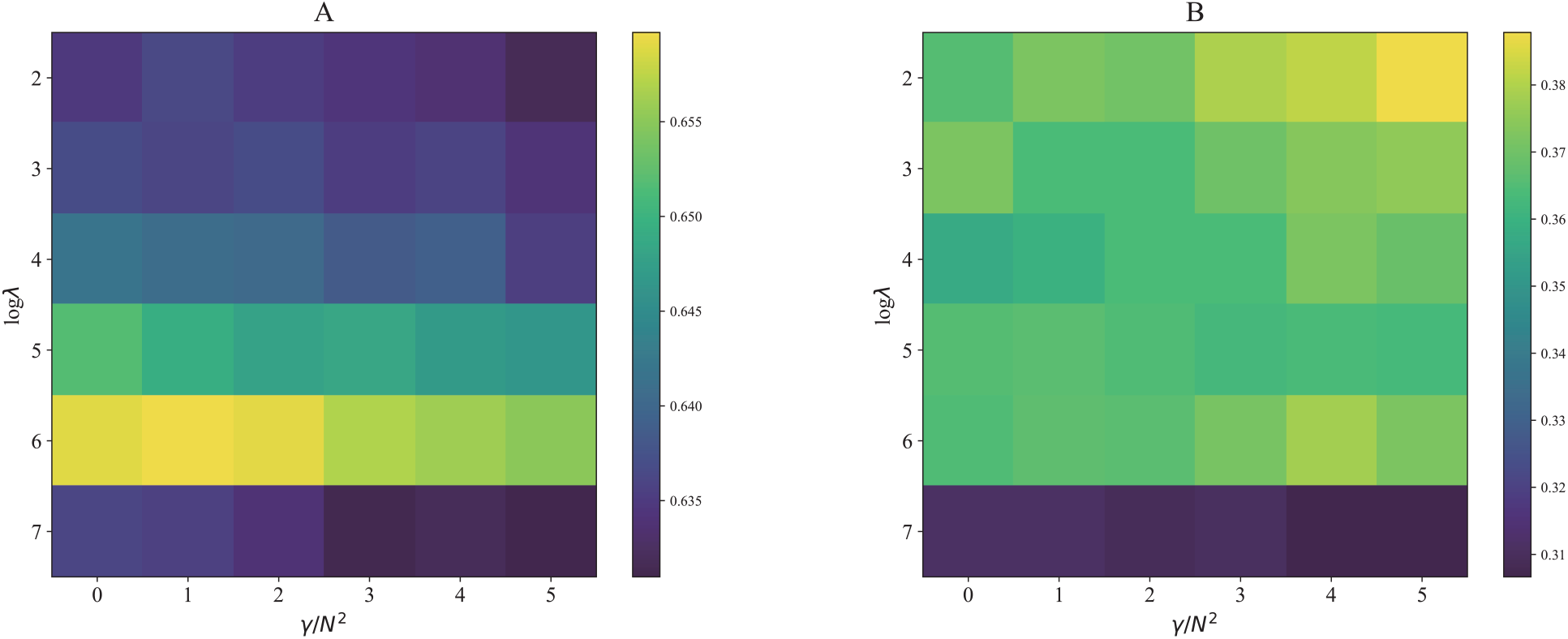
The parameter sensitivity results of *λ* and *γ* in terms of (**A**) ACC and (**B**) FRAC.

**Fig. 4.**
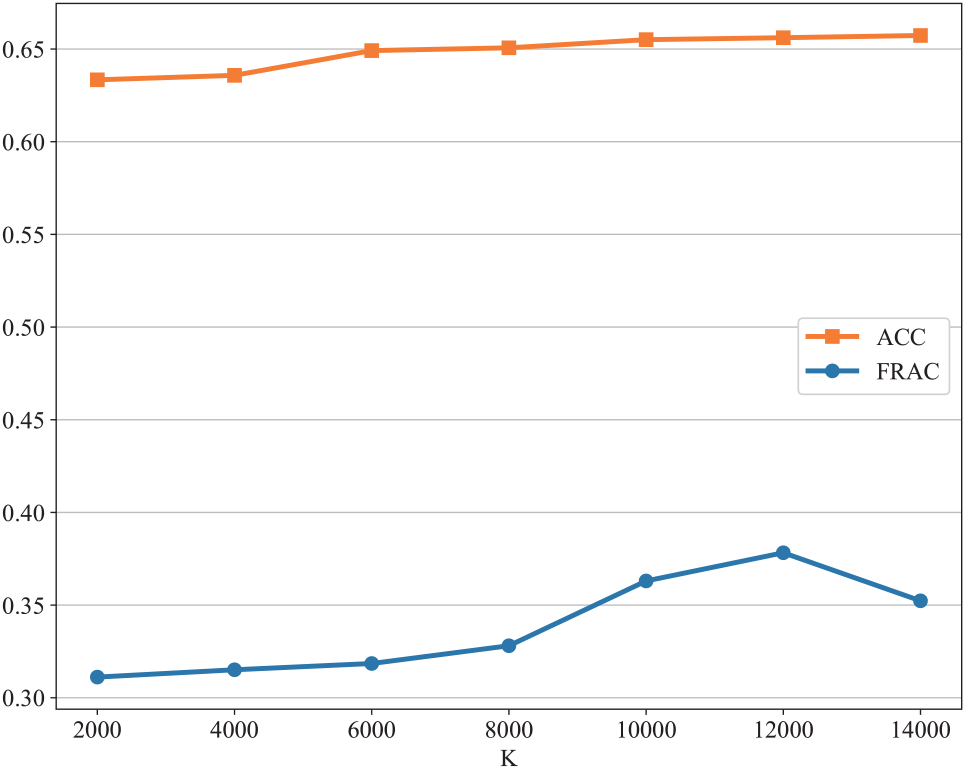
The parameter sensitivity results of *K*.

### 3.4. Case study

To illustrate the effectiveness of our model in identifying the common and unique protein complexes in different states, we introduce two partially overlapping protein complexes identified by PS-SNC from the Sign-293T network and Sign-HCT116 network.

Figures 5A and 5B show the sub-networks in Sign-293T network and Sign-HCT116 network, respectively, from which PS-SNC detected two partially shared protein complexes, i.e., complexes I and II. In particular, complex I contains the 293T cell-specific protein CDH2 [24]. We also check the most significantly enriched GO term for each predicted complexes using the web service of GO Term Finder (http://go.princeton.edu/cgi-bin/GOTermFinder). Complex I is enriched with GO term (GO:0045216) cell-cell junction organization. Complex II is enriched with GO term (GO:0030178) negative regulation of Wnt signaling pathway. Note that HCTT116 cells are colorectal carcinoma-derived cells. Colorectal cancer is one of the common malignancies worldwide and the Wnt signaling pathway is recognized as the main disrupted pathway in this malignancy [42, 43]. Therefore, PS-SNC can detect biologically meaningful complexes in different states effectively. Using PS-SNC to cluster the PPI networks of normal cells and cancer cells jointly can detect protein complexes related to cancer-induced mutations, which helps to explain the pathogenesis.

**Fig. 5.**
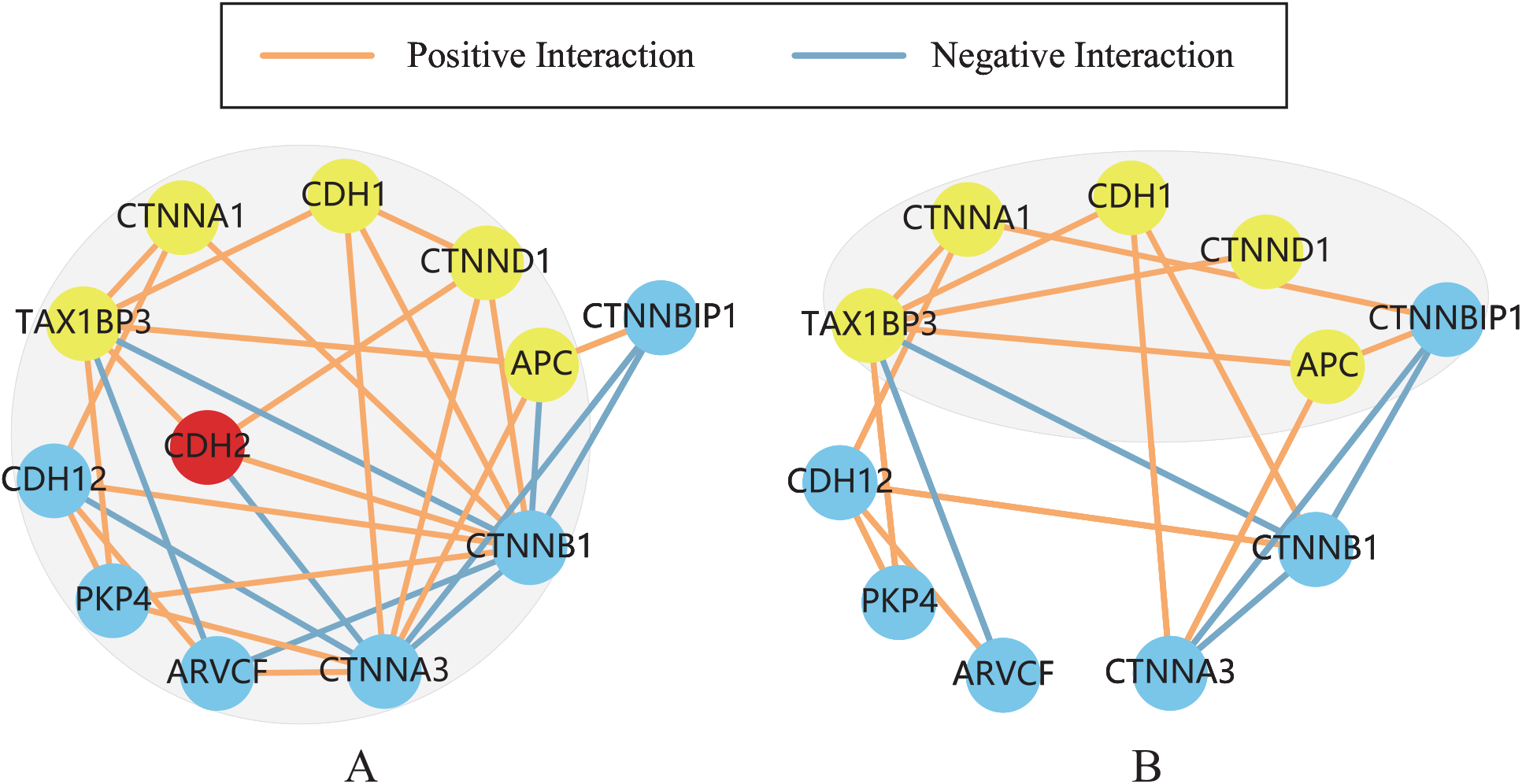
(**A**) The predicted complex I in Sign-293T network. (**B**) The predicted complex II in Sign-HCT116 network. The shadow areas show the predicted complexes I and II in Sign-293T network and Sign-HCT116 network respectively, yellow nodes represent the proteins shared across the two predicted complexes, red nodes represent proteins specific to the Sign-293T network, and blue nodes represent other proteins.

## 4. Conclusion

In this study, we propose a novel multi-network clustering model, named PS-SNC, to discover protein complexes from two signed PPI networks jointly. Our model can not only utilize the sign information of PPIs to guide the identification of protein complexes, but also explore the common and unique protein complexes in different networks. Extensive experimental results on synthetic and real datasets show that PS-SNC can improve the accuracy of protein complex identification effectively, and provide new insights for understanding the underlying mechanisms of disease and cell cycle developments. Furthermore, our model is a flexible framework which can be easily extended to the joint clustering of multiple signed networks.

## Funding

This work is supported by the National Natural Science Foundation of China [62173235, 61602309], Guangdong Basic and Applied Basic Research Foundation [2019A1515011384, 2022A1515010146], and the (Key) Project of Department of Education of Guangdong Province [No. 2022ZDZX1022].

## Notes

### Competing Interest Statement

The authors have declared no competing interest.

## References

[1] Bo Li and Bo Liao. Protein complexes prediction method based on core—attachment structure and functional annotations. International journal of molecular sciences, 18(9):1910, 2017.

[2] Daniel P Bondeson, Brenton R Paolella, Adhana Asfaw, Michael V Rothberg, Thomas A Skipper, Carly Langan, Gabriel Mesa, Alfredo Gonzalez, Lauren E Surface, Kentaro Ito, et al. Phosphate dysregulation via the xpr1–kidins220 protein complex is a therapeutic vulnerability in ovarian cancer. Nature Cancer, pages 1–15, 2022.

[3] Oron Vanunu, Oded Magger, Eytan Ruppin, Tomer Shlomi, and Roded Sharan. Associating genes and protein complexes with disease via network propagation. PLoS computational biology, 6(1):e1000641, 2010.

[4] Liang Yu, Jianbin Huang, Zhixin Ma, Jing Zhang, Yapeng Zou, and Lin Gao. Inferring drug-disease associations based on known protein complexes. BMC medical genomics, 8(2):1–13, 2015.

[5] Anne-Claude Gavin, Patrick Aloy, Paola Grandi, Roland Krause, Markus Boesche, Martina Marzioch, Christina Rau, Lars Juhl Jensen, Sonja Bastuck, Birgit Dümpelfeld, et al. Proteome survey reveals modularity of the yeast cell machinery. Nature, 440(7084):631–636, 2006.

[6] Nevan J Krogan, Gerard Cagney, Haiyuan Yu, Gouqing Zhong, Xinghua Guo, Alexandr Ignatchenko, Joyce Li, Shuye Pu, Nira Datta, Aaron P Tikuisis, et al. Global landscape of protein complexes in the yeast saccharomyces cerevisiae. Nature, 440(7084):637–643, 2006.

[7] Takashi Ito, Tomoko Chiba, Ritsuko Ozawa, Mikio Yoshida, Masahira Hattori, and Yoshiyuki Sakaki. A comprehensive two-hybrid analysis to explore the yeast protein interactome. Proceedings of the National Academy of Sciences, 98(8):4569–4574, 2001.

[8] Peter Uetz, Loic Giot, Gerard Cagney, Traci A Mansfield, Richard S Judson, James R Knight, Daniel Lockshon, Vaibhav Narayan, Maithreyan Srinivasan, Pascale Pochart, et al. A comprehensive analysis of protein–protein interactions in saccharomyces cerevisiae. Nature, 403(6770):623–627, 2000.

[9] Anne-Claude Gavin, Markus Bösche, Roland Krause, Paola Grandi, Martina Marzioch, Andreas Bauer, Jörg Schultz, Jens M Rick, Anne-Marie Michon, Cristina-Maria Cruciat, et al. Functional organization of the yeast proteome by systematic analysis of protein complexes. Nature, 415(6868):141–147, 2002.

[10] Trevor Clancy and Eivind Hovig. From proteomes to complexomes in the era of systems biology. Proteomics, 14(1):24–41, 2014.

[11] Anton J Enright, Stijn Van Dongen, and Christos A Ouzounis. An efficient algorithm for large-scale detection of protein families. Nucleic acids research, 30(7):1575–1584, 2002.

[12] Gary D Bader and Christopher WV Hogue. An automated method for finding molecular complexes in large protein interaction networks. BMC bioinformatics, 4(1):1–27, 2003.

[13] Min Li, Jian-er Chen, Jian-xin Wang, Bin Hu, and Gang Chen. Modifying the dpclus algorithm for identifying protein complexes based on new topological structures. BMC bioinformatics, 9(1):1–16, 2008.

[14] Min Wu, Xiaoli Li, Chee-Keong Kwoh, and See-Kiong Ng. A core-attachment based method to detect protein complexes in ppi networks. BMC bioinformatics, 10(1):1–16, 2009.

[15] Osamu Maruyama and Ayaka Chihara. Nwe: Node-weighted expansion for protein complex prediction using random walk distances. In Proteome science, volume 9, pages 1–11. BioMed Central, 2011.

[16] Tamás Nepusz, Haiyuan Yu, and Alberto Paccanaro. Detecting overlapping protein complexes in protein-protein interaction networks. Nature methods, 9(5):471–472, 2012.

[17] Sriganesh Srihari and Hon Wai Leong. A survey of computational methods for protein complex prediction from protein interaction networks. Journal of bioinformatics and computational biology, 11(02):1230002, 2013.

[18] Nazar Zaki, Dmitry Efimov, and Jose Berengueres. Protein complex detection using interaction reliability assessment and weighted clustering coefficient. BMC bioinformatics, 14(1):1–9, 2013.

[19] Eileen Marie Hanna and Nazar Zaki. Detecting protein complexes in protein interaction networks using a ranking algorithm with a refined merging procedure. BMC bioinformatics, 15(1):1–11, 2014.

[20] Marco Pellegrini, Miriam Baglioni, and Filippo Geraci. Protein complex prediction for large protein protein interaction networks with the core&peel method. BMC bioinformatics, 17(12):37–58, 2016.

[21] Shu Tadaka and Kengo Kinoshita. Ncmine: Core-peripheral based functional module detection using near-clique mining. Bioinformatics, 32(22):3454–3460, 2016.

[22] Zhourun Wu, Qing Liao, and Bin Liu. A comprehensive review and evaluation of computational methods for identifying protein complexes from protein–protein interaction networks. Briefings in bioinformatics, 21(5):1531–1548, 2020.

[23] Arunachalam Vinayagam, Jonathan Zirin, Charles Roesel, Yanhui Hu, Bahar Yilmazel, Anastasia A Samsonova, Ralph A Neumüller, Stephanie E Mohr, and Norbert Perrimon. Integrating protein-protein interaction networks with phenotypes reveals signs of interactions. Nature methods, 11(1):94–99, 2014.

[24] Edward L Huttlin, Raphael J Bruckner, Jose Navarrete-Perea, Joe R Cannon, Kurt Baltier, Fana Gebreab, Melanie P Gygi, Alexandra Thornock, Gabriela Zarraga, Stanley Tam, et al. Dual proteome-scale networks reveal cell-specific remodeling of the human interactome. Cell, 184(11):3022–3040, 2021.

[25] Le Ou-Yang, Dao-Qing Dai, and Xiao-Fei Zhang. Detecting protein complexes from signed protein-protein interaction networks. IEEE/ACM transactions on computational biology and bioinformatics, 12(6):1333–1344, 2015.

[26] Le Ou-Yang, Xiao-Fei Zhang, Dao-Qing Dai, Meng-Yun Wu, Yuan Zhu, Zhiyong Liu, and Hong Yan. Protein complex detection based on partially shared multi-view clustering. BMC bioinformatics, 17(1):1–15, 2016.

[27] Le Ou-Yang, Hong Yan, and Xiao-Fei Zhang. A multi-network clustering method for detecting protein complexes from multiple heterogeneous networks. BMC bioinformatics, 18(13):23–34, 2017.

[28] Chris Soon Heng Tan, Ka Diam Go, Xavier Bisteau, Lingyun Dai, Chern Han Yong, Nayana Prabhu, Mert Burak Ozturk, Yan Ting Lim, Lekshmy Sreekumar, Johan Lengqvist, et al. Thermal proximity coaggregation for system-wide profiling of protein complex dynamics in cells. Science, 359(6380):1170–1177, 2018.

[29] Kai Bartkowiak and Klaus Pantel. A shocking protein complex. Nature, 538(7625):322–323, 2016.

[30] Daniel Lee and H Sebastian Seung. Algorithms for non-negative matrix factorization. Advances in neural information processing systems, 13, 2000.

[31] Jing Liu, Yu Jiang, Zechao Li, Zhi-Hua Zhou, and Hanqing Lu. Partially shared latent factor learning with multiview data. IEEE transactions on neural networks and learning systems, 26(6):1233–1246, 2014.

[32] Yuri Pritykin and Mona Singh. Simple topological features reflect dynamics and modularity in protein interaction networks. PLoS computational biology, 9(10):e1003243, 2013.

[33] Deng Cai, Xiaofei He, Jiawei Han, and Thomas S Huang. Graph regularized nonnegative matrix factorization for data representation. IEEE transactions on pattern analysis and machine intelligence, 33(8):1548–1560, 2010.

[34] Donglin Niu, Jennifer G Dy, and Michael I Jordan. Iterative discovery of multiple alternativeclustering views. IEEE transactions on pattern analysis and machine intelligence, 36(7):1340–1353, 2013.

[35] Xiaochun Cao, Changqing Zhang, Huazhu Fu, Si Liu, and Hua Zhang. Diversity-induced multi-view subspace clustering. In Proceedings of the IEEE conference on computer vision and pattern recognition, pages 586–594, 2015.

[36] Daniel D Lee and H Sebastian Seung. Learning the parts of objects by non-negative matrix factorization. Nature, 401(6755):788–791, 1999.

[37] Zhipeng Xie, Chee Keong Kwoh, Xiao-Li Li, and Min Wu. Construction of co-complex score matrix for protein complex prediction from ap-ms data. Bioinformatics, 27(13):i159–i166, 2011.

[38] Zhourun Wu, Qing Liao, and Bin Liu. idenpc-miip: identify protein complexes from weighted ppi networks using mutual important interacting partner relation. Briefings in Bioinformatics, 22(2):1972–1983, 2021.

[39] Madalina Giurgiu, Julian Reinhard, Barbara Brauner, Irmtraud Dunger-Kaltenbach, Gisela Fobo, Goar Frishman, Corinna Montrone, and Andreas Ruepp. Corum: the comprehensive resource of mammalian protein complexes—2019. Nucleic acids research, 47(D1):D559–D563, 2019.

[40] Zhourun Wu, Qing Liao, Shixi Fan, and Bin Liu. idenpc-cap: Identify protein complexes from weighted rna-protein heterogeneous interaction networks using co-assemble partner relation. Briefings in Bioinformatics, 22(4):bbaa372, 2021.

[41] Le Ou-Yang, Min Wu, Xiao-Fei Zhang, Dao-Qing Dai, Xiao-Li Li, and Hong Yan. A two-layer integration framework for protein complex detection. BMC bioinformatics, 17(1):1–14, 2016.

[42] Meisam Jafarzadeh and Bahram M Soltani. Mirna-wnt signaling regulatory network in colorectal cancer. Journal of Biochemical and Molecular Toxicology, 35(10):e22883, 2021.

[43] Barbara Lustig, Boris Jerchow, Martin Sachs, Sigrid Weiler, Torsten Pietsch, Uwe Karsten, Marc van de Wetering, Hans Clevers, Peter M Schlag, Walter Birchmeier, et al. Negative feedback loop of wnt signaling through upregulation of conductin/axin2 in colorectal and liver tumors. Molecular and cellular biology, 22(4):1184–1193, 2002.

